# Genomic variant identification methods alter *Mycobacterium tuberculosis* transmission inference

**DOI:** 10.1101/733642

**Authors:** Katharine S. Walter, Caroline Colijn, Ted Cohen, Barun Mathema, Qingyun Liu, Jolene Bowers, David M. Engelthaler, Apurva Narechania, Julio Croda, Jason R. Andrews

## Abstract

Pathogen genomic data are increasingly used to characterize global and local transmission patterns of important human pathogens and to inform public health interventions. Yet there is no current consensus on how to measure genomic variation. We investigated the effects of variant identification approaches on transmission inferences for *M. tuberculosis* by comparing variants identified by five different groups in the same sequence data from a clonal outbreak. We then measured the performance of commonly used variant calling approaches in recovering variation in a simulated tuberculosis outbreak and tested the effect of applying increasingly stringent filters on transmission inferences and phylogenies. We found that variant calling approaches used by different groups do not recover consistent sets of variants, often leading to conflicting transmission inferences. Further, performance in recovering true outbreak variation varied widely across approaches. Finally, stringent filters rapidly eroded the accuracy of transmission inferences and quality of phylogenies reconstructed from outbreak variation. We conclude that measurements of genetic distance and phylogenetic structure are dependent on variant calling approach. Variant calling algorithms trained upon true sequence data outperform other approaches and enable inclusion of repetitive regions typically excluded from genomic epidemiology studies, maximizing the information gleaned from outbreak genomes.

## Introduction

The continuous evolution of human pathogens creates a powerful epidemiological record. Patterns of variation within and between populations of pathogens can be used to infer substitution rates, phylogenetic and phylogeographic relationships, such as geographic origins and routes of spatial spread, population size dynamics, and – if pathogen evolution occurs over the same timescale as transmission – transmission patterns^1^.

Tuberculosis (TB) kills more people than any other infectious disease and halting transmission of *Mycobacterium tuberculosis* is essential to reducing the global burden of disease. However, in high-incidence settings, it is unknown where and between whom the majority of transmission occurs^2–4^ and therefore where to focus interventions. Patterns of *M. tuberculosis* genetic and genomic variation are frequently used to identify potential recent transmission events. *M. tuberculosis* isolates that share a genotype (RFLP, spoligotype, or MIRU-VNTR)^5–7^, or whose whole genome sequences are within a given genetic distance^8–11^, are considered clustered and potentially epidemiologically linked. Phylogenies inferred from outbreak variation may reveal patterns of relatedness within and between clusters^11–13^. Finally, transmission trees integrate epidemiological and phylogenetic information to capture probable transmission histories, chains of who infected whom^14,15^. Predicted transmission links have been used to infer the likely location and/or timing^16,17^ of transmission, to identify risk factors for transmission and high risk populations^18^, to distinguish between acquired (primary) and transmitted drug resistance^19^, and to declare an outbreak over^20^.

Transmission inferences in molecular epidemiology for *M. tuberculosis* and other pathogens rely on the high-quality measurement of genetic variation from sequence data. However, there is no consensus on how to measure pathogen genomic variation, and studies frequently employ different sequence quality control measures, mapping algorithms, variant callers, and variant filters^21^. Further, the performance of variant calling methods for different pathogen species is not well described, meaning that uncertainty in the underlying genotypic or sequence data used to inform transmission inferences is unmeasured. The *ad hoc* nature of genomic variant calling makes it difficult to interpret pathogen variation identified within a study and to compare variation across studies. Whether variant calling methods affect transmission inferences and the accuracy of different methods in measuring variation within outbreaks has not been assessed.

The lack of standardized approaches for pathogen genomic epidemiology results in part from the fact that many genomic tools and approaches have been designed and validated for the measurement of human genomic variation^21^. Commonly used variant callers may call diploid genotypes (i.e. VarScan^22^, DeepVariant^23^), be trained on human sequence data (i.e. DeepVariant^23^) or require truth sets of known segregating mutations to train models (i.e. GATK’s VQSR^24^) and variant truth sets are required to measure performance of variant calling pipelines. Variant calling pipelines optimized for human genomes likely perform differently on pathogen genomes, which differ significantly in within-species diversity and genomic characteristics including ploidy, G-C content, genomic architecture, and repetitive content. A recent study found that commonly used variant calling pipelines have poor accuracy^25^ when applied to bacterial genomes, highlighting the importance of benchmarking variant callers for bacterial species.

Variant calling methods may perform differently upon different species and, additionally, for different applications. Standard workflows for *M. tuberculosis* genomic epidemiology generate short-read sequence data from cultured isolates and then filter variants both by region and variant annotations^26^. While many pipelines widely used in *M. tuberculosis* molecular epidemiology were designed or validated for antibiotic resistance prediction^27–32^, their performance in recovering true pairwise differences and the underlying phylogenetic structure of outbreak genomes, the metrics used for transmission inference, has not been reported.

Here, we investigate the effects of variant calling approaches on transmission inference of *M. tuberculosis*. First, to assess variability of commonly used and published pipelines in the tuberculosis genomic epidemiology field, we collected and compared variant calls from five research groups for the same sequence data from a clonal outbreak in Germany^33^. Second, we measured the performance of variant calling tool combinations in recovering genome-wide variants and pairwise differences between outbreak genomes in a simulated tuberculosis outbreak for which we knew the underlying genomic truth. Finally, we identify general aspects of variant identification approaches that can improve transmission inference.

## Methods A. Reanalyzing a clonal *M. tuberculosis* outbreak

Molecular epidemiology studies harness genetic and genomic variation to understand patterns of transmission and are premised on the idea that the *M. tuberculosis* is constantly evolving as it spreads from person to person. Currently, the standard approach in *M. tuberculosis* genomic epidemiology studies is to sequence whole genomes directly from bacterial cultures, generating short-read Illumina sequence data^26,34^. Sequence data are mapped to a reference genome with one of several widely used mapping algorithms (including BWA, Bowtie2, and SMALT), variants are identified with respect to the reference genome using a variant calling algorithm (including GATK and Samtools), and variants are filtered, often by applying hard filters that specify thresholds for variant annotations, such as variant quality score or depth.

We tested the effect of variant calling approaches on phylogenetic and transmission inference by collecting variants from four molecular epidemiology groups (A-D) for previously published sequence data. We invited two groups with published variant calling pipelines specific for *M. tuberculosis* (B and C), using the variant caller GATK, and two groups using Samtools, another widely used variant caller, in order to explore the effects of applying different variant pipelines recently published in the *M. tuberculosis* genomic epidemiology literature. We sought to test the effect of applying different pipelines to the same genomic data; this comparison was not intended to be an exhaustive comparison of published pipelines. We additionally included the original set of published variant calls (E)^33^, for a total of five pipelines for comparison. Pipeline characteristics are in Table S1.

Each group submitted variants they identified in sequence data from a clonal *M. tuberculosis* outbreak in Germany from 1997 – 2006^33^. The outbreak was identified during routine population-based surveillance and 86 isolates were cultured on Lowenstein Jensen Media and sequenced on an Illumina platform (ENA Study Accession: PRJEB6945). These published sequencing data are of varying quality; we used these data for comparison because, uniquely, they are accompanied by a partial “truth set” of Sanger-confirmed SNPs and because this dataset has been used for other transmission studies^14^.

### Pipelines

Groups submitted filtered variant calls as single-sample or multi-sample VCFs in addition to a multiple sequence alignment of concatenated single nucleotide polymorphisms (SNPs). We used LiftoverVcf (http://broadinstitute.github.io/picard/) to convert variant coordinates for pipelines A and B to coordinates on the H37Rv reference genome so that variant positions could be compared.

Pipeline C made diploid calls and did not provide a multiple sequence alignment. To create a multiple-sequence alignment of consensus sequences, we converted diploid calls to haploid by setting homozygous genotypes (0/0 or 1/1) to the corresponding haploid genotype (0 or 1) and heterozygous genotypes to the genotype with greater allele depth. We used bcftools^35^ to generate consensus sequences, setting genotypes that were absent in single-sample VCFs to the reference allele, as recommended by the authors. We then used snp-sites^36^ to select variants internal to the outbreak. This conservatively excludes minority variants. Transmission inferences using pairwise differences are most frequently made with the consensus genomes of individual isolates (although some studies use minority variants for transmission inferences^37^), and comparing minority variants was outside the scope of this study.

Pipeline A additionally included diploid calls, however, also provided a FASTA used for pairwise differences and phylogenies in addition to a list of variant sites internal to the outbreak. We used the list of variant sites internal to the outbreak to compare variant sites with other pipelines.

### Sensitivity

The true outbreak sequences are unknown and therefore the performance of variant calling pipelines in recovering true variants cannot be measured; however, the original study confirmed 85 single nucleotide polymorphisms with Sanger sequencing. We report pipelines’ sensitivity in recovering these high-confidence previously identified SNPs as a partial measure of sensitivity. Specificity could not be measured.

### Phylogenetic inference

We calculated raw pairwise differences between isolates with the R package ape v.5.2 (model = ‘ N’). We constructed maximum likelihood phylogenies with RAxML-ng^38^ with a GTR substitution model. We used a Stamatakis ascertainment bias correction to correct for invariant sites and specified nucleotide stationary frequencies present in the H37Rv genome. We measured phylogenetic distances between a random selection of 100 bootstrap replicate trees derived from SNPs from each pipeline using the Robinson-Foulds metric^39^. We reduced the dimensionality of tree distances with principal components analysis using the R package treespace^40^. We performed hierarchical clustering of trees using Ward’s method also in treespace^40^.

## Methods B. Measuring performance of variant calling tool combinations on simulated genomic data

Because the genomic truth for a true outbreak can never be known, we next simulated a tuberculosis outbreak and generated sequence data *in silico* (Fig. S1). We applied commonly used mapping and variant calling algorithms to simulated data and measured the performance of these variant calling tool combinations in recovering (1) true *M. tuberculosis* genomic variants and (2) true pairwise differences between closely related *M. tuberculosis* sequences. Here, our aim was to explore characteristics of the most accurate variant calling approaches rather than to compare pipelines of specific groups. We additionally tested how choice of reference genome effects performance by mapping variants against 12 different reference genomes of varying distance to the CDC1551 query genome (Table S2). Software versions are in Table S3.

### Simulated sequence data

We generated 20 independent Illumina readsets (2 × 151-bp) from the CDC1551 genome *in silico*, with the next generation sequence read simulator ART v. 2.5.8^41^, which simulates reads from a given genome with read lengths and error profiles from commonly used sequence platforms (Fig. S1). We simulated reads using a built-in quality profile for a HiSeqX v2.5 TruSeq sequencing machine. Before simulations, we set ambiguous sites in the CDC1551 genome to N. We simulated reads with a mean of 100X coverage, with a mean and standard deviation fragment length of 650-bp and 150-bp, respectively (consistent with Illumina recommended insert sizes of 350-bp (https://support.illumina.com/sequencing/sequencing_instruments/hiseq-x/questions.html; standard deviation from empirical data).

Measuring performance requires a truth VCF of true variant sites in the query genome with respect to a given reference genome (Fig. S1). To generate truth VCFs for the CDC1551 query genome with respect to 12 *M. tuberculosis* reference genomes (Table S2), we pairwise aligned the query genome (CDC1551) to each reference genome with MUMmer^42^ (*nucmer* with the *maxmatch* option). We identified SNP variants from the pairwise alignments using MUMmer *show-snps*, excluding SNPs with ambiguous mapping and indels.

### Mapping and variant calling

We mapped simulated reads with commonly used mapping algorithms (BWA, Bowtie 2, and SMALT), with default settings, to 12 reference genomes spanning described *M. tuberculosis* diversity and a distance of 1037 to 2901 SNPs from CDC1551 (Table S4). We called variants with commonly used variant callers (GATK and Samtools), setting ploidy to 1. We called variants for each sample independently rather than jointly calling genotypes because joint variant calling approaches are designed for human cohort studies and were found to be less sensitive in detecting singleton and low-frequency variants in a previous study^43^. Variant calling tools and versions are listed in Table S5.

We additionally called variants for each sample independently with DeepVariant v.0.7.0, a convolutional neural network trained upon human genomic truth sets to identify variants in short-read sequence data^23^. Specifically, DeepVariant v.0.7.0 was trained upon labeled genotypes from a total of 16 sets of human genomic data, including 10 PCR-free sequence replicates of HG001, 2 PCR-free replicates of HG005 PCR-free, and 4 PCR replicates of HG001. The genomic “truth” which the model is trained on includes variants that have been identified by several pipelines and occur within “high confidence” regions of the human genome^44^. The model was frozen after training and then can be applied to unseen genomic data in the form of aligned reads (BAM files). DeepVariant does not have an option to infer haploid genotypes; therefore, we assigned homozygous genotype predictions to the corresponding haploid call (i.e. assigning 0/0 to 0 and 1/1 to 1). For heterozygous calls, we used allele depth to assign genotype as the allele with greater coverage. If two alleles at a heterozygous site had equal depth, we randomly selected a haploid genotype. We set DeepVariant SNPs filtered as “RefCall” to missing. For all callers, we output all-sites VCF files (i.e. both variant and invariant sites) in order to distinguish between reference allele calls and missing or “no-call.”

### Quality filtering

We excluded indels and applied two independent filters to SNP variant calls to samples individually: (1) a single hard variant quality score filter, QUAL < 40 and (2) VQSR (variant quality score recalibration)^24^. VQSR fits Gaussian mixture models to annotations characterizing a truth set of high-quality variants and then applies this model to all candidate variants to recalibrate variant quality scores. Because a high-quality truth set does not exist for *M. tuberculosis*, we defined our truth set internally, including all candidate SNPs with a QUAL score greater than the mean QUAL score for a given set of variants. We set a phred-scaled prior likelihood of 15 and used the annotations DP, QD, MQRankSum, ReadPosRankSum, FS, SOR, and MQ in the model. We set the recalibrated variant quality score (VQSLOD) threshold so that our caller would have 99% sensitivity for recovering variants within our truth set. We did not apply VQSR to DeepVariant calls to avoid overfitting.

We filtered per site and per sample so that when making pairwise comparisons, a site that did not pass filters for a single sample was considered missing for that sample alone and could be included in comparisons between other samples.

### Performance benchmarking

We used hap.py^45^, software widely used to measure performance of variant calling pipelines upon human genomic variation^23^, to assess the performance of each pipeline in recovering true SNPs across the genome, within the 168 PE/PPE genes^46^, and outside of the PE/PPE genes.

### Outbreak simulations

To measure the performance of pipelines in recovering true pairwise differences between closely related samples, such as those sampled in an outbreak setting, we simulated a short, relatively densely-sampled tuberculosis outbreak with TransPhylo^14,47^. We simulated an outbreak that began in 2013 and was observed until 2018 with a basic reproduction number, R_0_, of 3 (Fig. S3). We set generation time, the time between subsequent infections, and sampling time, time between infection and diagnosis, as Gamma distributed, with shape = 10 and scale = 0.1, corresponding to a mean of one year. We set the product of the within-host population size and generation time (N_e_g) to 100/365 and the probability of observing cases, π, to 0.25.

TransPhylo simulates transmission trees, graphs of who infected whom and when in an outbreak. We extracted the underlying phylogeny from the simulated transmission tree. Using a substitution rate estimate of 2 substitutions/ site/ year, which falls within the range reported by a recent meta-analysis of the *M. tuberculosis* molecular clock^48^, we rescaled the simulated phylogeny, where branch lengths were in units of years, to a phylogeny with branch lengths in units of substitutions per site. We chose a rate on the higher end of published clock rates for *M. tuberculosis* to ensure that we would obtain sufficient numbers of “true” simulated SNPs on which to test both pipelines and downstream inference.

To generate a set whole genome sequences related by the simulated genealogy, we then simulated evolution along the simulated phylogeny with Pyvolve^49^. Pyvolve takes a phylogeny, a root sequence, and a nucleotide substitution model and simulates evolution along the branches of a phylogeny. We simulated nucleotide evolution from the CDC1551 reference genome with an F81 model of nucleotide evolution^50^ with empirically-derived nucleotide frequencies. We used snp-sites^36^ to generate a VCF file of variant sites in the tip genomes. Pyvolve introduces variants randomly along the root sequence; to simulate variation at sites known to be polymorphic in *M. tuberculosis*, we replaced the sites simulated with Pyvolve with randomly selected sites that varied between CDC1551 and H37Rv, allowing us to preserve the simulated phylogenetic structure while including variants that are segregating in natural *M. tuberculosis* populations. We applied this set of SNPs to the CDC1551 reference genome, generating 44 simulated outbreak sequences. We used LiftOver to generate a “truth” outbreak VCF with respect to the H37rV genome.

From each tip genome, we simulated Illumina short-read sequence data, mapped reads, and called variants as described above. We called variants for each sample individually and applied filters described above to individual sample variant files. We measured precision and recall in variant calling pipelines in detecting true pairwise SNP differences between simulated genomic sequences and measured the true and measured raw pairwise genetic distances (number of nucleotide differences between sequences) between samples with the R package *ape*^51^, using the dist.dna function.

### Phylogenetic inference

To determine the effect of filtering on phylogenetic inference, we focused on variants identified by a single tool combination, BWA/GATK. We then selected variant quality thresholds that corresponded to variant deciles, generated multiple alignments of SNPs meeting quality thresholds, and inferred maximum likelihood phylogenies for each multiple alignment. We fit maximum likelihood trees with RAxML-ng, with a GTR substitution model. We applied a Stamatakis ascertainment bias correction to correct for invariant sites and specified nucleotide stationary frequencies present in the CDC1551 outbreak root genome. We measured phylogenetic distances from the best supported trees to the true tree using the Robinson-Foulds distance^39^ and the Kendall-Colijn metric^52^, with lambda equal to 0.

## Results A

To measure the effect of variant calling pipeline on transmission inference, four groups (A-D) contributed filtered variant calls for previously published sequence data from a clonal *M. tuberculosis* outbreak in Hamburg and Schleswig-Holstein, Germany from 1997 – 2006^33^. The outbreak was identified during routine population-based surveillance and 86 isolates were cultured and fully sequenced on an Illumina platform (ENA Study Accession: PRJEB6945). The original study identified 85 SNPs that were validated with Sanger sequencing^33^.

Variant calling pipelines varied in quality control, choice of reference genome, mapper, caller, variant filters, and genomic regions excluded (Table S1). In this section, we aim to highlight the effect of applying different pipelines to the same data rather than to isolate the effect of any single component of a pipeline on variants identified (Results B).

### Variant calling pipelines identify different sets of *M. tuberculosis* genomic variants when applied to the same sequence data

After filtering, pipelines identified 63 to 416 SNPs between outbreak strains (i.e. internal SNPs) compared to 85 SNPs identified in the initial study (Fig. 1a, Table S2). The five pipelines identified a common set of 55 SNPs (Fig. 1b); however, there was significant discordance in SNPs identified and each pipeline identified 1-190 unique SNPs. Sensitivity in recovering SNPs confirmed by Sanger sequencing in the original study ranged from 72.9 – 92.9% (Fig. 1c, Table 1). Two variants identified by pipeline B fell in locations on the pipeline B’s reference genome (one of the outbreak genomes) that did not correspond to references used by other groups and therefore were unique due to reference choice. Pipeline C excluded 20% (17/86) of samples that did not meet thresholds for contamination (minimum of 90% of reads taxonomically classified as *M. tuberculosis* complex)^29^.

**Figure 1.**
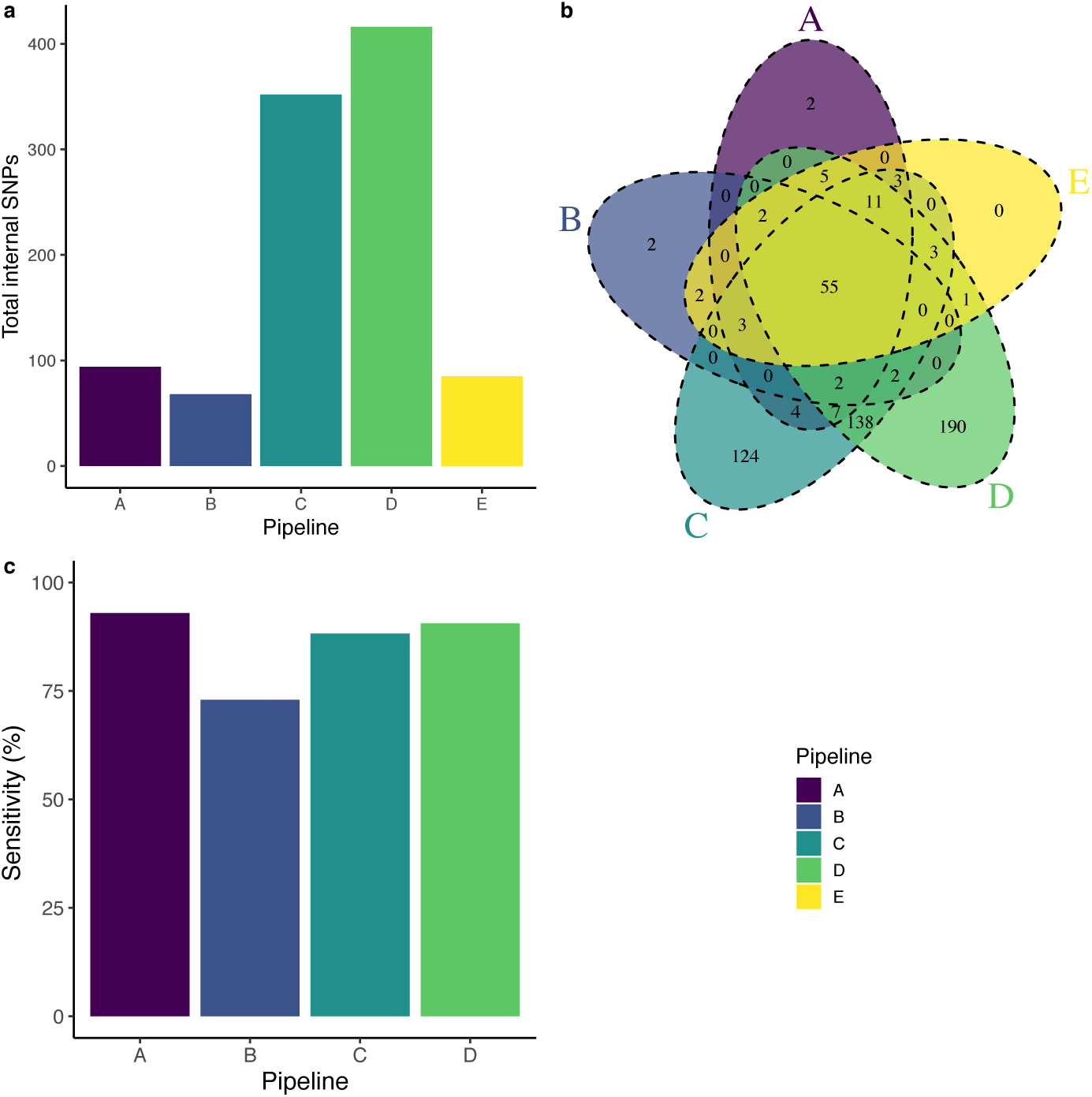
Different pipelines call different variants when applied to the same sequence data. (a) Total internal SNPs identified by each pipeline (A-D) compared with the 85 SNPs detected in the original study (E). (b) Venn diagram of the intersection of SNPs identified by each pipeline. (c) Sensitivity of each pipeline (A-D) in recovering the set of Sanger sequence-verified SNPs from the original study (E).

Importantly, the high number of variants identified by pipelines C and D is likely in part a result of pipelines generating VCF files that include only variant sites. This results in no information about non-variant sites in the final variant files, making it difficult to distinguish between sites with a confident reference allele call and sites with no confident allele call (i.e. at positions of low coverage or quality). Often, the assumption is made that non-variant sites are indeed the reference allele, resulting in inflated measures of pairwise differences.

### Differing variant calls result in different transmission inferences

Pairwise genetic distances between outbreak sequences, a proxy of the evolutionary distance between genomes, are frequently used to identify *M. tuberculosis* isolates potentially linked by recent transmission^53^. Two isolates separated by a large evolutionary distance are considered unlikely to be the result of recent transmission, while isolates within a threshold genetic distance^8–11^ are considered clustered and potentially epidemiologically linked. Public Health England, for example, prioritizes clusters for further “targeted public health investigation and action.”^8^ While distance thresholds vary across studies^53,54^, 5^9^- or 12-SNP^8^ thresholds are frequently used to distinguish between “clustered” and “non-clustered” isolates^26^. To test the effect of variant calling pipeline on predicted transmission, we applied these two distance thresholds.

The five pipelines identified different distributions of pairwise SNP distances (Fig. 2a), corresponding to widely different epidemiological interpretations (Fig. 2a,b). Median pairwise distances ranged from 1 to 42 SNPs among pipelines (Table S2). 0 – 29.7% of isolate pairs were identical (0 SNP differences). After applying commonly used transmission thresholds of pairwise distances less than or equal to 5 or 12 SNPs^8,28,55^, the number of potential transmission links varied dramatically across pipelines (Fig. 2b, Table S2). For example, by pipeline A, 80.7% of sample comparisons fell below a 5-SNP threshold of potential recent transmission whereas by pipelines C and D, less than 0.5% of comparisons did.

**Figure 2.**
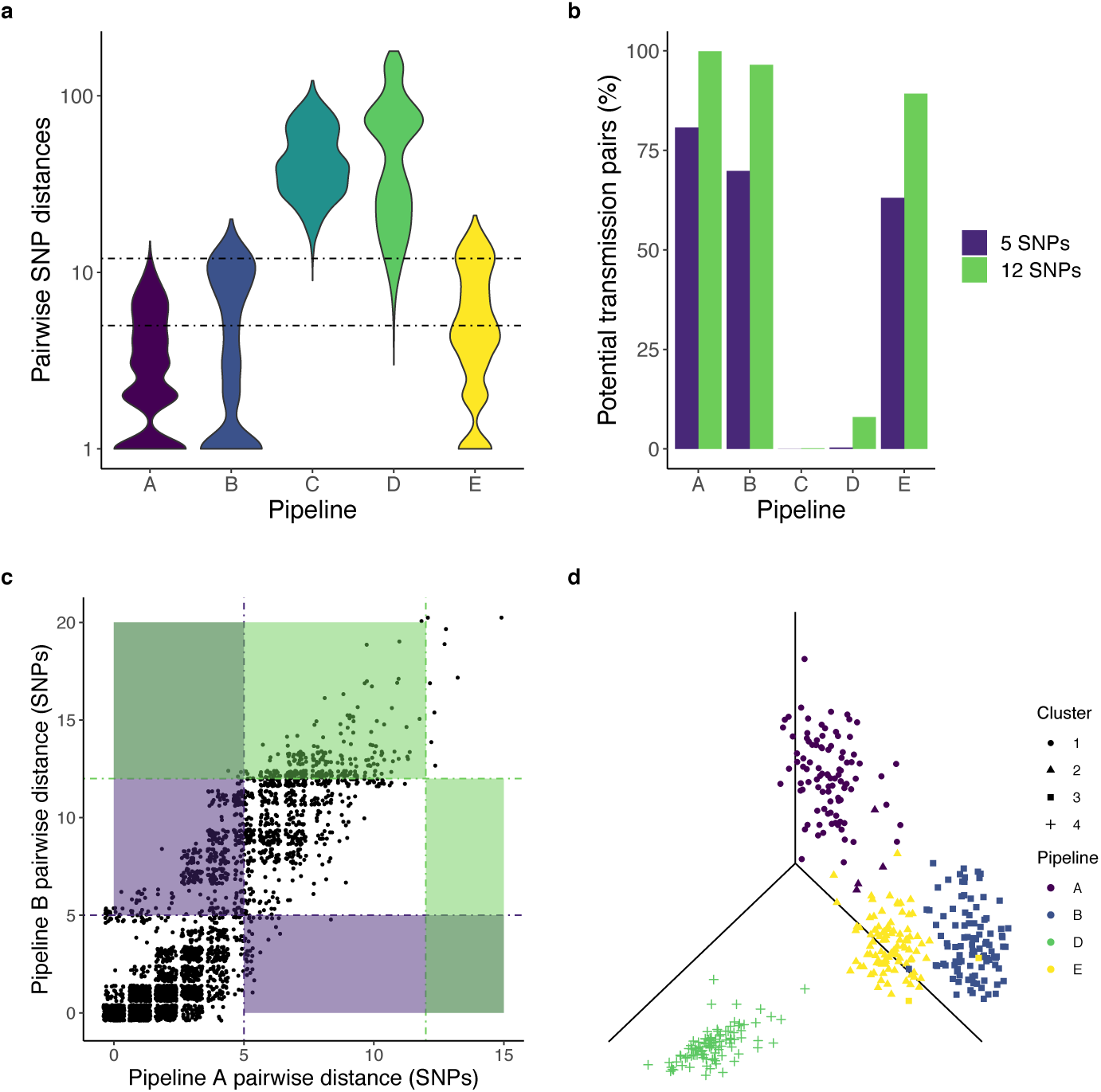
Pairwise SNP distances and phylogenies differ across variant calling pipelines. (a) Violin plot of the distribution of pairwise genetic distances identified by each pipeline on a log-scale. The width of the violin represents the frequency of a given pairwise genetic distance. The dotted lines at 5 and 12 SNPs represent commonly used thresholds for recent transmission, above which, the possibility of recent transmission is excluded. Pipelines A, B, D, and E include 86 sequences (3655 comparisons). Pipeline C includes 69 sequences passing quality filters (2346 comparisons). (b) The percentage of sequence pairs with potential transmission links when applying 5 and 12 SNP thresholds for transmission. (c) Pairwise distances between isolates identified by pipelines A and B. Each point corresponds to a unique pair of sequences. Dotted lines indicate 5 and 12 SNP distance thresholds, commonly used for inferring recent transmission. Blue and red shading indicates regions in which callers make conflicting transmission inferences after applying a 5 or 12 SNP threshold respectively. (d) Maximum likelihood trees inferred from the variation identified by each pipeline largely cluster separately. Tree distances between 100 bootstrap replicate trees from each pipeline were measured with the Robinson-Foulds (RF) metric and summarized by principal components analysis. The first three axes are shown; color indicates variant calling pipeline and shape indicates the tree cluster assigned with Ward’s method. Because RF distances require trees to have identical tips, trees from pipeline C, which only included calls for 68 samples, are not included.

Even pipelines that identify similar total numbers of internal SNPs (A and B) and that identify pairwise differences that are closely correlated (Fig. 2c, r = 0.89, p < 0.001) may still identify different distances between isolate pairs (Fig. 2c), resulting in conflicting transmission inferences. After applying a 5-SNP threshold for transmission, Pipeline A identifies 413 potential clustered pairs not identified by pipeline B. Conversely, pipeline B identifies 14 potential clustered pairs not identified by pipeline A. Cumulatively, for the two most similar pipelines, 11.7% (427/ 3655) of transmission inferences are discordant (Fig. 2c, blue shading). For all other pipeline comparisons, discordance was substantially greater.

### Differing variant calls result in different phylogenetic inferences

To test the effect of pipelines on phylogenies, we fit maximum likelihood phylogenies with alignments of concatenated SNPs identified by each pipeline. We assessed the similarity of bootstrapped trees with Robinson Foulds distance and used Ward’s method to assign trees into clusters (Fig. 2d). Trees inferred from variants identified by different pipelines are largely assigned to distinct clusters (Fig. 2d). However, 4% (4/100) bootstrap replicate trees constructed with Pipeline A variants cluster with Pipeline E trees and 2% (2/100) bootstrap replicate trees constructed with Pipeline E variants cluster with Pipeline B trees. (Pipeline C cannot be compared and is therefore not shown because tree distances cannot be computed between trees with different sets of tips.)

## Results B

For the outbreak described above, as for any outbreak, the true genomic sequence of *M. tuberculosis* isolates is unknown. Performance of pipelines in recovering true outbreak SNPs cannot be measured. Variant calling pipelines for human genomes are often benchmarked upon diploid human genomic “truth sets,” variants identified and confirmed by several sequencing and bioinformatic pipelines and/or validated by family pedigrees^56,57^. However, such genomic variant truth sets do not exist for *M. tuberculosis* or other human pathogens.

To measure the performance of commonly used variant calling tool combinations in identifying (a) *M. tuberculosis* SNPs and (b) identifying pairwise distances between closely related isolates, we simulated *M. tuberculosis* short-read sequence data from complete, published *M. tuberculosis* genomes (Fig. S1). We applied commonly used mapping algorithms (BWA, Bowtie 2, and SMALT), variant callers (GATK, Samtools, and DeepVariant), and filters (a hard quality score filter, QUAL, and Variant Quality Score Recalibration, VQSR) to simulated data (Methods) and measured performance in recovering true SNP variants (Methods). Approaches used in genomic epidemiology studies vary widely in choice of mapper, caller, and filters. We explore only a subset of possible tool combinations here.

### Performance in recovering true *M. tuberculosis* SNPs varies across tool combinations and reference genomes

To measure performance of variant calling tools in recovering genome-wide *M. tuberculosis* variants, we generated 20 sets of Illumina short-read data *in silico* from the CDC1551 query genome and evaluated the performance of nine variant calling tool combinations in recovering the true 1107 SNPs between the query and the frequently used H37Rv reference genome (Fig. S1).

Performance in recovering true genome-wide *M. tuberculosis* SNPs varies widely across tool combinations (Fig. 3) using H37Rv as the mapping reference. Prior to filtering, variation in precision exceeds that of recall; maximum precision is 83.7% (BWA/GATK) while maximum recall is 94.7% (SMALT/DeepVariant) (Table S3). The F1 score, the harmonic mean of precision (positive predictive value) and recall (sensitivity), commonly used to rank genomic pipelines^23^, varies from 0.791 (Bowtie2/DeepVariant) to 0.886 (BWA/GATK) (Table S3) before filtering.

**Figure 3.**
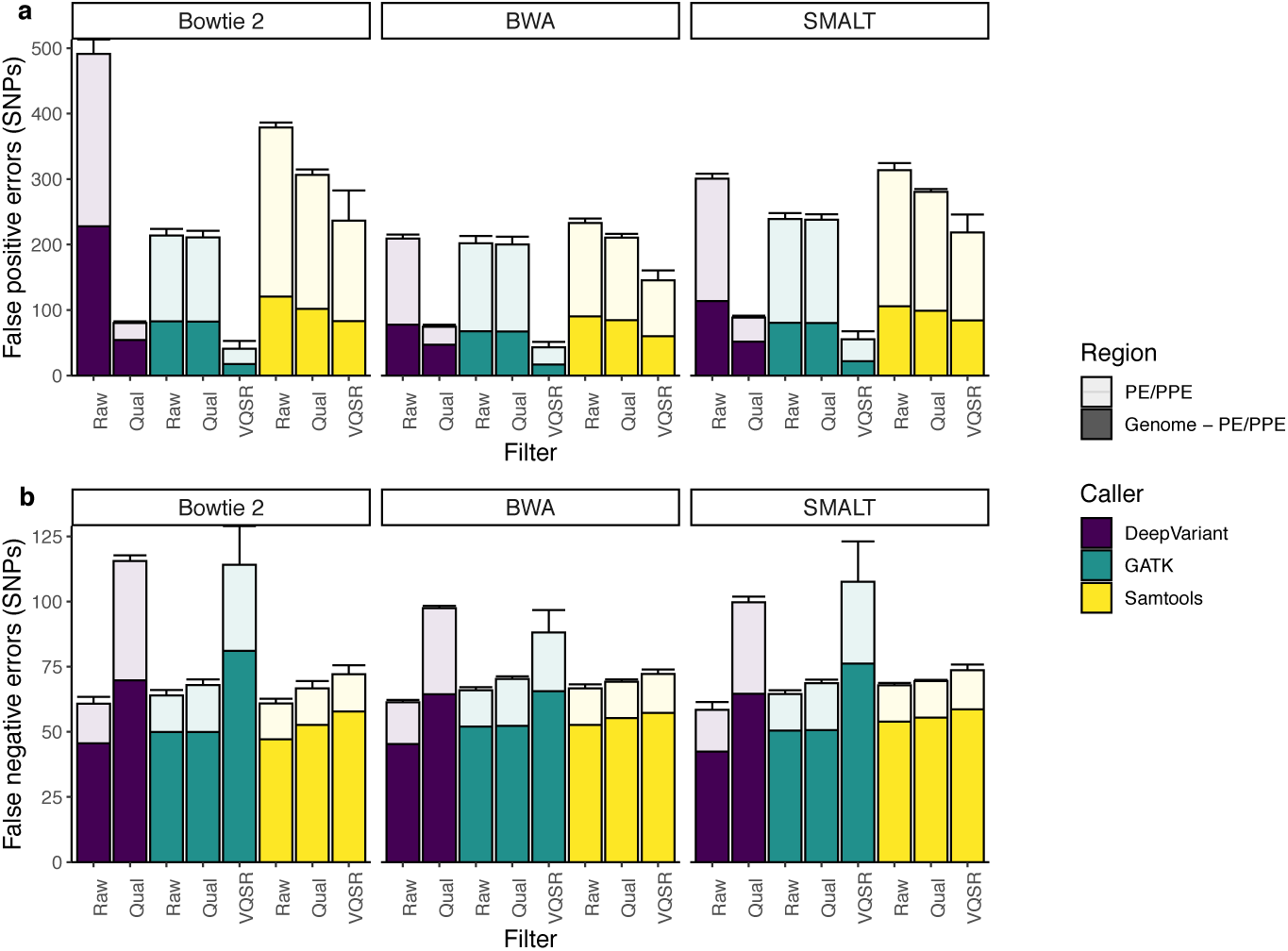
False positive and false negative errors in *M. tuberculosis* SNPs vary across tool combinations. Mean false positive (a) and false negative (b) errors identified by the nine tool combinations of read mapper and variant caller and three filters. Filters include Raw, no filtering; Qual, excluding variants with low quality score (QUAL < 40); and VQSR, variant quality score recalibration (VQSR only applied to GATK and Samtools calls) (Methods). Bar color indicates mapper and bar shading indicates genomic region: light shading indicates errors falling within the 168 PE/PPE genes and dark shading indicates errors falling outside that region. Bars and error bars indicate the mean and standard deviation of 20 replicates. Panels have different y-axes.

We examined the genomic location of errors and tested if standard filters could reduce FP errors. Variant calling performance varies across the genome and is worse in the 168 repetitive PE/PPE genes^46^, which are often excluded from *M. tuberculosis* molecular epidemiology studies (Fig. 3). Before filtering, 53.6 – 68.2% (identified by Bowtie 2/DeepVariant and Bowtie 2/Samtools, respectively) of FPs occur in PE/PPE genes, which comprise 6.37% of the genome (Fig. 3a). FN errors are also disproportionately located in the PE/PPE genes, though to a lesser extent (Fig. 3b). Before filtering, 20.5 – 27.4% (identified by (SMALT/Samtools and SMALT/DeepVariant, respectively) of FNs occur in PE/PPE genes.

Filtering by excluding the PE/PPE genes or by filtering by quality score or VQSR reduces but does not eliminate FP errors, while increasing FN errors. FP errors are minimized by BWA/GATK with VQSR and excluding the PE/PPE genes (mean FP = 16.8 SNPs, mean FN = 65.6 SNPs). Even when PE/PPE genes are included, GATK/VQSR tool combinations identify fewer FP errors than all other tool combinations (Fig. 3). FN errors are minimized by SMALT/DeepVariant excluding the PE/PPE genes (mean FP = 114 SNPs, mean FN =42.4 SNPs).

All tool combinations are characterized by a trade-off between recall and precision visible in the inverse relationship between false positive (FP) and false negative (FN) errors. While the maximum F1 score was 95.5% (BWA/GATK/VQSR, excluding PE/PPE genes), no tool combination consistently outperforms other tool combinations in minimizing both types of errors (Fig. 3), indicating that optimal approach may depend on the relative costs of different error types for specific applications. No combination of mapper, variant caller, and filter were able to achieve >99.9% precision and recall reported for human genomes and which won the PrecisionFDA Truth Challenge^23^.

### Performance in recovering true *M. tuberculosis* SNPs improves with closely related reference genomes

To assess how choice of reference genome effects variant calling performance, we mapped 20 replicate sequence sets to 12 different reference genomes spanning global *M. tuberculosis* diversity and ranging from 1037 (Lineage 4, strain F11) to 2901 (Lineage 2, strain Beijing_NITR203) SNPs distant from the CDC1551 query genome (Lineage 4, Table S4). In a general linear model, log-transformed distance to the reference genome, mapper, and caller are significant predictors of FP errors and log-transformed distance to the reference genome and caller are significant predictors of FN errors prior to filtering.

Both FP and FN errors increase with increasing log-transformed distance between the query and reference genomes, when controlling for mapper and caller (FP errors: r = 0.47, p-value < 0.001; FN errors: r = 0.43, p-value < 0.001) (Fig. S2). However, errors vary widely between reference genomes, possibly reflecting individual genomes’ repetitive content, extent of synteny with the query genome, or reference assembly quality. Interestingly, the F1 score was slightly positively correlated with distance to the reference genome (Fig. S2). This results from the negative correlation between recall and log-transformed distance from the reference genome (r = −0.16, p-value < 0.001) and the positive correlation between precision and distance from the reference genome (r = 0.36, p-value < 0.001).

### Performance in recovering true pairwise differences between outbreak strains varies across tool combinations

The goal of genome wide variant calling – used to identify genomic correlates of antibiotic resistance or virulence, for example – is to identify mutations between a single query genome and a reference genome. In contrast, variant calling for transmission inference seeks to measure small amounts of variation between multiple closely related outbreak genomes. Identifying variants between query genomes and a known reference genome is intermediate to the true goal: identifying variants between the outbreak genomes. If errors with respect to the reference genome are consistent within a single pipeline, then inference about relatedness between outbreak samples should not be affected.

To measure the performance of tool combinations in identifying pairwise differences between closely related sequences, we simulated a five-year tuberculosis outbreak (Methods). We simulated evolution of *M. tuberculosis* from a common ancestral genome (CDC1551) over the outbreak phylogeny (Fig. S3), resulting in a total of 147 SNPs internal to the outbreak, and generated sequence data *in silico* from the 44 outbreak sequences. True pairwise differences between outbreak genomes ranged from 0-27 SNPs and mean pairwise distances between isolates was 13.4 SNPs.

Performance in recovering true pairwise differences between outbreak strains varied across tool combinations using H37Rv as the mapping reference. Prior to filtering, mean recall ranges from 91.0% (BWA/Samtools) to 93.5% (Bowtie2/DeepVariant) and mean precision ranges from 14.7% (Bowtie 2/Samtools) to 64.5% (SMALT/GATK). As seen for genome-wide performance, performance in measuring pairwise differences is worse in the 168 repetitive PE/PPE genes compared to the rest of the genome. Before filtering, 40.8% (BWA/DeepVariant) – 97.0% (BWA/GATK) of FPs occur in PE/PPE genes (Fig. 4).

**Figure 4.**
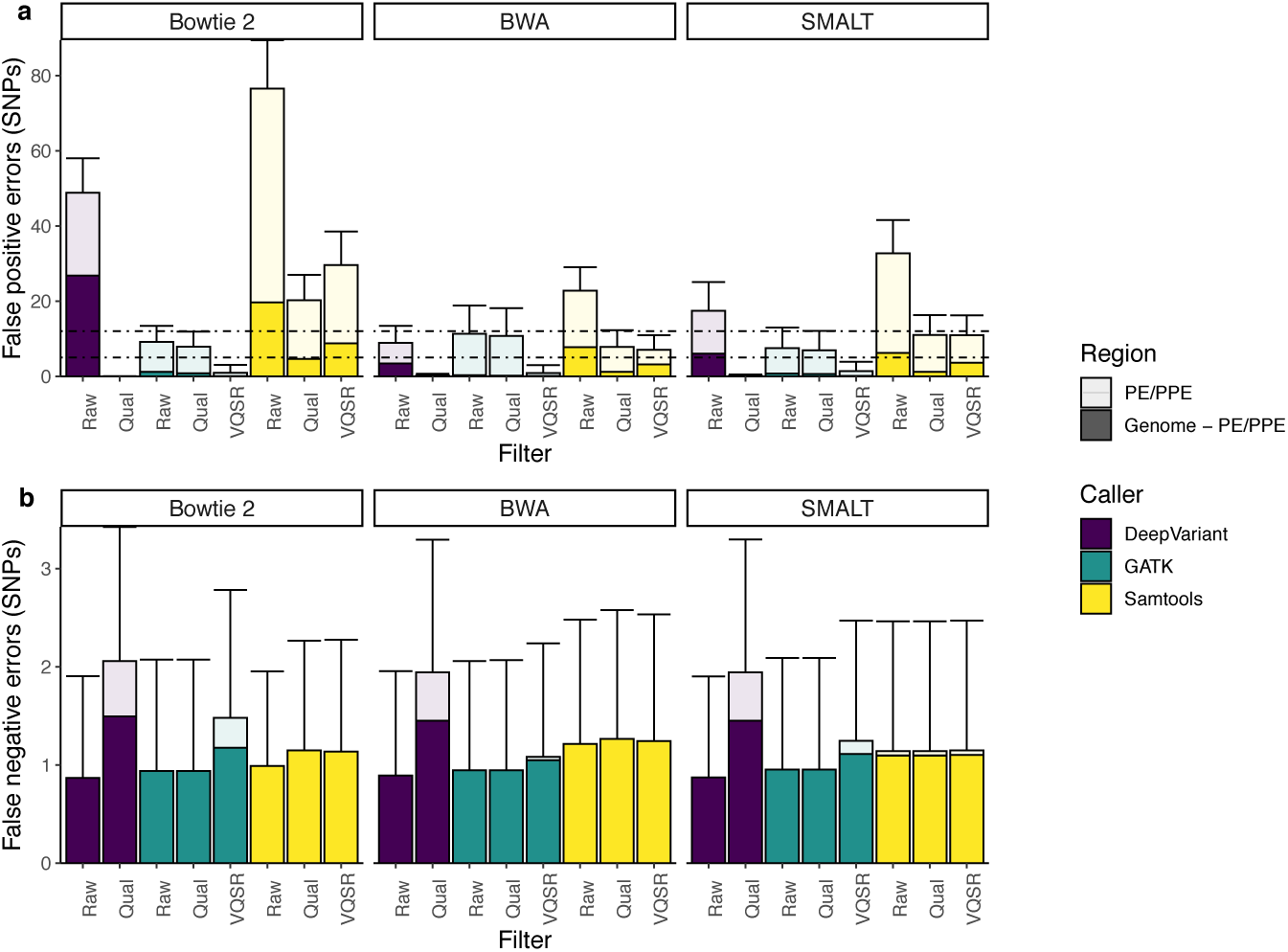
False positive and false negative errors in pairwise differences between *M. tuberculosis* genomes vary across tool combinations. Mean false positive (a) and false negative (b) SNP differences for each pairwise comparison of the 44 outbreak sequences (946 pairwise comparisons) identified by the nine tool combinations of read mapper and variant caller and three filters. Filters include Raw, no filtering; Qual, excluding variants with low quality score (QUAL < 40); and VQSR, variant quality score recalibration (VQSR only applied to GATK and Samtools calls) (Methods). Bar color indicates mapper and shading indicates genomic region: light shading indicates errors falling within the 168 PE/PPE genes and dark shading indicates errors falling outside that region. Bars indicate the mean across pairwise comparisons; error bars extend to one standard deviation above the mean. Dotted lines in (a) indicate 5 and 12 SNP distance thresholds, commonly used for inferring recent transmission. Panels have different y-axes.

We then tested whether filters could improve performance in recovering pairwise differences. Quality score or VQSR filters reduce but do not eliminate pairwise errors. Even after filtering, the range of mean FP errors is more than 26 times that of FN errors across tool combinations (Fig. 4). Several tool combinations result in mean FP pairwise errors above 5 SNPs if PE/PPE genes are not excluded (i.e. approaches with Samtools and with GATK/QUAL). If a pairwise difference threshold of 5 SNPs was applied, the effect of variant calling errors alone would exclude the possibility of recent transmission.

Because 23 of the 147 randomly introduced outbreak SNPs occur within PE/PPE genes, approaches which exclude PE/PPE genes have a maximum total sensitivity of only 84.4% (124/147) of the total outbreak variation. Tool combinations including GATK/VQSR or DeepVariant/QUAL allow PE/PPE gene variation to be retained while keeping mean FP errors below 5 SNPs. Among tool combinations which include PE/PPE gene variation and with mean FP errors below 5 SNPs, maximum recall was 91.6% (BWA/GATK/VQSR) and maximum precision was 96.3% (Bowtie2/DeepVariant/QUAL).

We investigated the source of persistent FP errors – errors outside PE/PPE genes that were not eliminated by filters – in two of the best-performing tool combinations, BWA/GATK/VQSR and BWA/DeepVariant/QUAL. Five of 7 positions with persistent FP errors after VQSR occurred at positions that had failed a VQSR filter for other samples, though not the query samples, indicating that sites which fail VQSR for a single sample are potentially problematic sites that should be excluded for all samples. For BWA/DeepVariant/QUAL calls, 2 of 3 persistent FP pairwise SNPs were at positions that were initially called as heterozygous by the diploid variant caller and the third was due to lack of coverage at a site in resulting in erroneous reference allele calls. This suggests that haploid variant calling algorithms and further training of variant calling algorithms on *M. tuberculosis* genomic variation could further improve variant calling accuracy.

### Increased variant filtering may hinder transmission inferences

Variant filters vary widely between studies and can contribute more to variation between tool combinations than either mapping or variant calling (Figs. 3 and 4). However, filters are frequently not justified empirically, and the effect of filtering on transmission and phylogenetic inference is unknown. To test the effect of variant filtering on downstream inferences, we generated ten sets of Illumina sequence data in silico for the outbreak samples described above and applied a series of increasingly stringent quality score filters to variant calls identified by a single tool combination, BWA/GATK. We used the distribution of all variant quality scores to identify quality score thresholds that demarcated deciles of variants so that we would exclude an additional 10% of identified variants with each increase of the quality score filter. For each set of increasingly filtered variants, we measured the accuracy of identifying isolate pairs falling within a 5-SNP threshold and fit maximum likelihood trees to SNP alignments. We measured distance of inferred trees to the underlying true outbreak tree using both the Kendall-Colijn metric (KC)^52^ and the Robinson-Foulds distance (RF)^39^.

As expected, applying increasingly strict variant quality score filters reduces observed pairwise differences between outbreak samples, resulting in a trade-off between FP and FN errors (Fig. 5a). Mean genome-wide FP pairwise errors are 10.5 SNPs before quality filtering and 0.29 SNPs after excluding the PE/PPE genes. Mean genome-wide FP errors fall rapidly to 0.14 after excluding variants in the lowest quality decile and 0.045 SNPs after excluding PE/PPE genes. Mean genome-wide FN errors are 0.95 before filtering and increase after excluding the lowest two deciles of variants. FN errors are consistently higher in variant sets excluding PE/PPE genes, reflecting the fact that 15.6% (23/147) of true variants occur in these genes.

**Figure 5.**
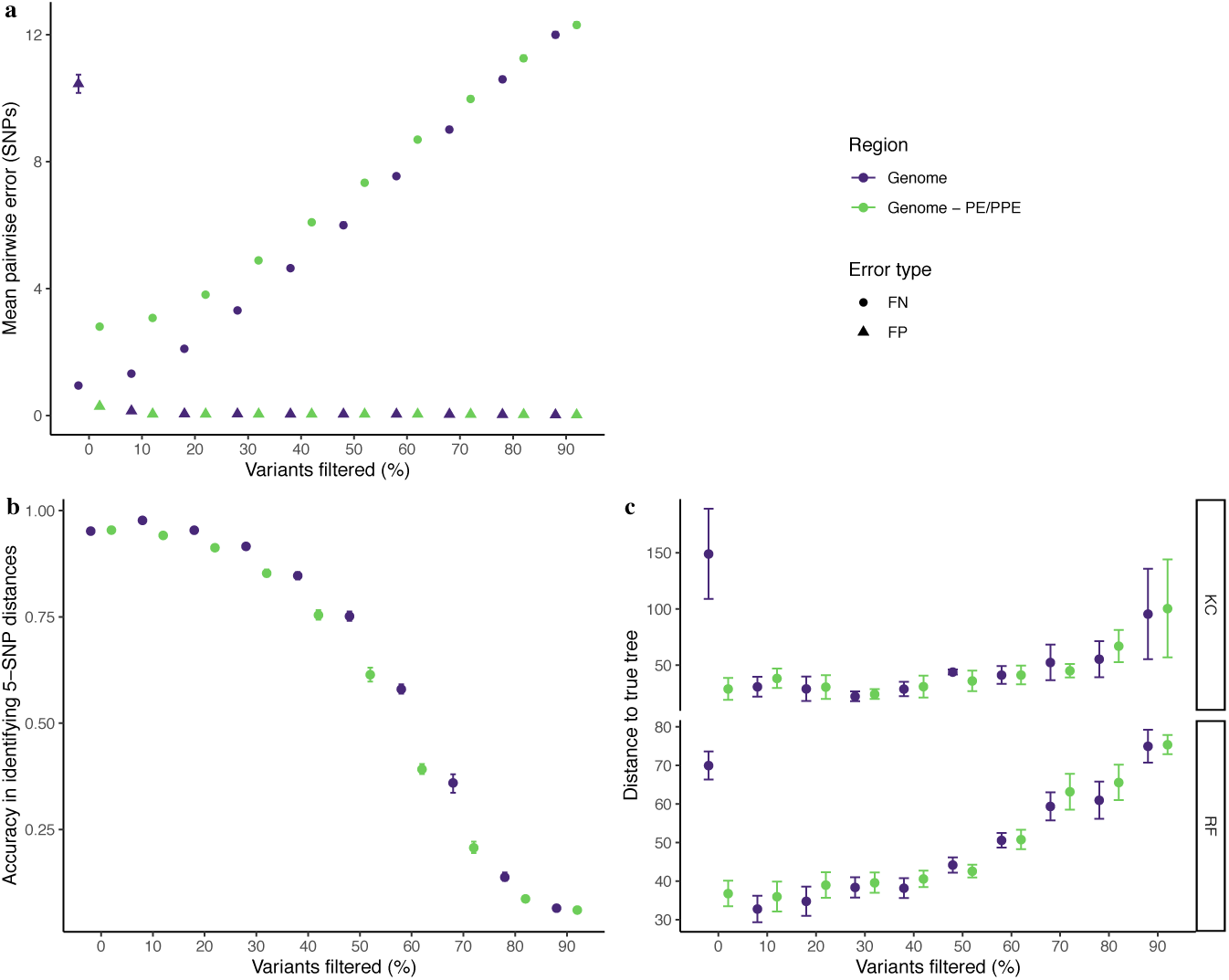
Increased filtering does not always improve transmission inference and phylogenetic reconstruction. Ten replicate Illumina sequence sets for the simulated tuberculosis outbreak were generated *in silico* and variants identified with BWA/GATK and filtered with increasing stringency (Methods). X-axes indicate the percentage of total variants excluded as an increasingly strict filtering approach was applied. Genomic region is indicated by color and points corresponding to each genomic region are staggered along the x-axis to improve clarity. (a) False positive pairwise errors decrease and false negative pairwise errors increase with increasingly strict variant filtering. Errors were identified by comparing the recovered variants with the underlying true variants. Error type is indicated by point shape and genomic region is indicated by color. Points represent mean FN and FP SNP errors and error bars indicate the partially pooled errors across ten replicate sequence sets. (b) Accuracy in distinguishing pairs falling above or below a 5-SNP threshold decreases with increasingly stringent filters. Error bars indicate the partially pooled errors across ten replicate sequence sets. (c) Distance of phylogenies inferred with increasingly filtered sets of variants to the true outbreak phylogeny. Panels indicate tree distance metric: KC, Kendall-Colijn metric^52^ and RF, Robinson-Fould’s distance^39^. Error bars indicate standard deviation distances for ten replicate sets of sequence data generated *in silico* for the simulated outbreak.

Before quality filtering, 95.2% of isolate pairs were correctly assigned as falling above or below a 5-SNP threshold when considering genome-wide variants; 95.4% pairs were correctly assigned after exclusion of the PE/PPE genes (Fig. 5b). Accuracy in distinguishing pairs falling above or below a 5-SNP threshold improves slightly after excluding variants in the lowest quality decile to a maximum of 97.7% for genome-wide variants and 94.2% after excluding PE/PPE genes, after which accuracy rapidly declines. Excluding the PE/PPE genes generally results in lower accuracy in identifying isolate pairs falling under a 5-SNP threshold (Fig. 5b).

Distances of reconstructed trees to the true underlying phylogeny fall rapidly after initial filtering and then steadily increase with more stringent quality score filters, resulting in a U-shaped relationship between quality score filter and distance to the true tree, measured by KC distance, and a hockey-stick shaped relationship for RF distance (Fig. 5c). When no quality filtering is applied, the inclusion of variants within PE/PPE genes results in large distances of inferred phylogenies to the true tree (149.0, KC distance and 70.0, RF distance). Mean KC tree distances fall to a minimum of 22.4 after filtering 30% of variants, when genome-wide variants are included. Mean RF distances fall to a minimum of 35.8 after filtering of 10% of variants, when genome-wide variants are included. These observations suggest that some filtering is necessary to remove the lowest quality variants, either by exclusion of problematic regions or by exclusion of the lowest quality variants, but additional filtering may rapidly erode the quality of inferred phylogenies. Further, regional filters and quality score filters interact to determine overall accuracy of inferences based on pairwise differences and phylogenies.

## Discussion

While many applications of *M. tuberculosis* whole-genome sequencing for transmission inference use hard filters to minimize false positive SNPs^26^ and then apply pairwise SNP distance thresholds to infer potential transmission linkages^26,34^, here we show that a) such approaches do not recover consistent sets of SNPs; b) pairwise distance thresholds are not robust to differences between pipelines; and c) strict filtering does not always improve transmission inferences made using pairwise differences or phylogenies.

As shown in part A, methodological differences between different *M. tuberculosis* molecular epidemiology groups can lead to differing epidemiological conclusions made from the same sequence data. This suggests that results from genomic epidemiology studies need to be interpreted in the context of study methodology. The degree of diversity or clonality reported within a single outbreak, for example, may reflect methodology rather than true outbreak diversity. Additionally, pairwise distance thresholds for recent transmission developed using one variant calling pipeline^55^ cannot be easily adapted by studies using different pipelines. Similarly, estimates of *M. tuberculosis* substitution rate are contingent on variant calling pipeline. For example, excluding variants within the PE/PPE genes discards the most variant-dense regions of the *M. tuberculosis* genome and will likely decrease the observed molecular clock rate of *M. tuberculosis*. For genomic information to be pooled across studies, a standardized variant calling approach needs to be applied as was recently done in a meta-analysis estimating *M. tuberculosis* substitution rates^48^. Further, differences in filters represent an important source of difference in variants identified.

Sequencing technologies and variant calling algorithms are rapidly changing, and our aim was not to identify a single best pipeline. However, we found that performance varies widely among approaches and identified several characteristics of variant calling approaches that may improve the accuracy of variant calling for transmission inference (Box 1). We found that variant calling performance metrics do not translate across species; tools developed and tested on human data perform measurably worse on *M. tuberculosis* compared to human genomes. The best performing pipelines included either DeepVariant with a quality score filter or GATK with VQSR, highlighting the benefit of variant caller calibration upon labeled sequence data, either through the training of a neural network or fitting of Gaussian mixture models to variant annotations.

### Box 1: General considerations for pathogen variant calling for transmission inference

- Apply a taxonomic filter and select reads corresponding to the taxa of interest or exclude reads mapping to other taxa^65^.
- Generate long sequence reads or paired-end reads, if using short-read data.
- Map reads to a closely related reference genome or assemble reads de novo to generate an outbreak reference genome.
- Output invariant and variant sites to distinguish between reference allele calls and positions without a confident allele call (potentially corresponding to deletions or regions with low- or poor-quality sequence coverage).
- Call variants for samples independently rather than with a joint variant calling approach^43^. (Joint variant calling approaches are designed for human cohort studies and previous studies found them less sensitive in detecting singleton and low-frequency variants^43^, classes of variants valuable for transmission inferences.)
- Apply variant callers calibrated upon sequence data, such as neural networks or callers with sophisticated error models fit to variant annotations.
- Conduct filtering on site and sample-specific annotations (i.e. many variant calling programs “merge” sample-specific annotations into a maximum or mean annotation for a site).
- Apply filters to both reference allele calls and alternate allele calls (i.e. reference allele calls might have poor coverage and/or quality just as alternate alleles might).
- Retain the greatest extent of the genome possible.
- Use a haploid-specific caller or convert genotypes to correct ploidy.
- Publish pipeline scripts and software, including versions.

As expected, variant errors increased with increased distances between the reference and query genomes. This contrasts with a previous study that found choice of reference genome did not affect *M. tuberculosis* epidemiological inferences^58^. Our study differs from the previous study in that we used simulated genomic data for which underlying true variation is known to measure performance in identifying variants in individual genomes. The earlier study measured how reference choice affects performance in classifying isolate pairs as linked or unlinked using transmission links identified using the CDC1551 reference genome as truth.

*M. tuberculosis* genomic epidemiology studies routinely use the H37Rv or CDC1551 reference genomes, both of which belong to Lineage 4. Studies investigating variation in other lineages will particularly benefit from using local reference genomes, either a full-length genome from the outbreak being studied or another closely related genome. Gene content differs between *M. tuberculosis* lineages^26,59^, constraining sensitivity in a reference-based genome approach. Any variation within regions inserted in the query genomes relative to the reference will be missed even by a perfectly sensitive variant caller. The use of a local reference genome by one of the pipelines in Part A, for example, enabled the identification of two variants unobserved by other groups because they occurred within regions inserted relative to the standard H37Rv reference. Generating longer reads and/or assembly of full-length pathogen genomes will further reduce the errors intrinsic to mapping-based approaches. The effect of reference choice is likely to be even more pronounced for other bacterial species with greater diversity.

Our finding that the cumulative effect of false positive and false negative errors in pairwise SNP differences frequently exceeds commonly used thresholds for recent transmission events suggests that molecular epidemiology studies need to be interpreted with caution. Subtle differences between outbreak genomes can be readily overwhelmed by variant errors. However, appropriate filtering greatly reduced both false positive and false negative errors while retaining variation in PE/PPE genes, indicating that errors are largely predictable and can be minimized with appropriate error models.

Many genomic epidemiology studies employ some type of hard filtering, whether based on annotation or genomic region. Transmission inferences based on pairwise differences as well as phylogenies are sensitive to variant filtering strategy and optimal filters may depend on specific downstream application. While minimal filtering improves the accuracy of transmission linkages predicted by pairwise differences and tree reconstruction, extensive filtering results in poorer accuracy of predicted transmission linkages and phylogenies that are increasingly distant from the underlying true phylogeny. After limited quality filtering, including the PE/PPE genes does not negatively affect transmission or phylogenetic inferences. The PE/PPE genes are the most variant dense regions of the *M. tuberculosis* genome and are known antigens and virulence determinants^46^. Routine exclusion of these genes reduces the information potential of *M. tuberculosis* genomes and limits our ability to study the functional consequences of *M. tuberculosis* variation. Further, we found that more sophisticated error models outperform hard annotation-based filters and identified tool combinations that minimize error while retaining the PE/PPE genes.

While our focus is on *M. tuberculosis*, a bacterium that is considered to be slow-evolving^48,55,60^, the issues we identify here generalize to other pathogens^61^. Our results suggest that pathogen genomic epidemiology, for *M. tuberculosis* and other species, will benefit from genomic resources similar to those that exist for human genomes. First, pathogen genomic truth sets, experimental (not simulated) sequence data accompanied by validated variants, would enable training of machine learning approaches upon labeled pathogen variant data and would serve as a gold standard for performance benchmarking of variant calling approaches. Secondly, further work is needed to optimize variant callers for pathogens and for particular applications (i.e. prediction of antibiotic resistance versus transmission inference). For example, variant callers could output quality scores for reference allele calls in addition to alternative allele calls, enabling comparisons between all sites (as GATK already does). Finally, variant uncertainty represents an important and unreported source of potential error in genomic epidemiology studies. How to incorporate uncertainty in underlying measures of genomic variants or sequences in phylogenetic inference remains an open and important question for genomic epidemiology and population genomics more broadly.

Here, we focused on the measurement of SNPs from short-read sequence data for transmission inference. Indels and other structural variants are an important additional source of *M. tuberculosis* variation that contain important phylogenetic information, which we did not examine here^62^. Nor do we examine within-host variation, which is clinically and epidemiologically important^63,64^ and can provide additional information for transmission inference^37^. Further, we measured performance on simulated sequence data, that generates sequences with platform-specific error profiles^41^, but which likely does not capture the full spectrum of sequence errors from epidemiological studies. In addition, we generated genomic “truth sets” by pairwise aligning query and reference genomes. The genomic truth therefore depends on the accuracy of this pairwise alignment. Additionally, we did not investigate the performance of variant calling tools in recovering variants associated with antibiotic resistance or other clinically and epidemiologically important phenotypes. Transmission inference is one of many potential applications of WGS data and the characteristics of variant callers optimal for transmission inference are not necessarily those optimal for resistance predictions.

Our findings demonstrate that current measures of pathogen genomic variation are susceptible to errors that may be propagated through all downstream analyses. Developing genomic resources and methods for specific pathogen species and harnessing the power of long-read sequence data will improve accuracy of transmission inferences and enable measurement of uncertainty in molecular epidemiology studies.

## Acknowledgements

We thank Matthew Ezewudo and Marco Schito for supplying variant calls from the UVP pipeline^29^, Caitlin Pepperell for comments, and John Burton Hanks for computational support. This work was supported by the National Institute of Allergy and Infectious Disease at the National Institutes of Health (grant R01 AI130058 to JRA) and the Stanford Child Health Research Institute (Postdoctoral Support Award to KSW).

