## Supplementary Information for "Genomic variant identification methods alter *Mycobacterium tuberculosis* transmission inference"

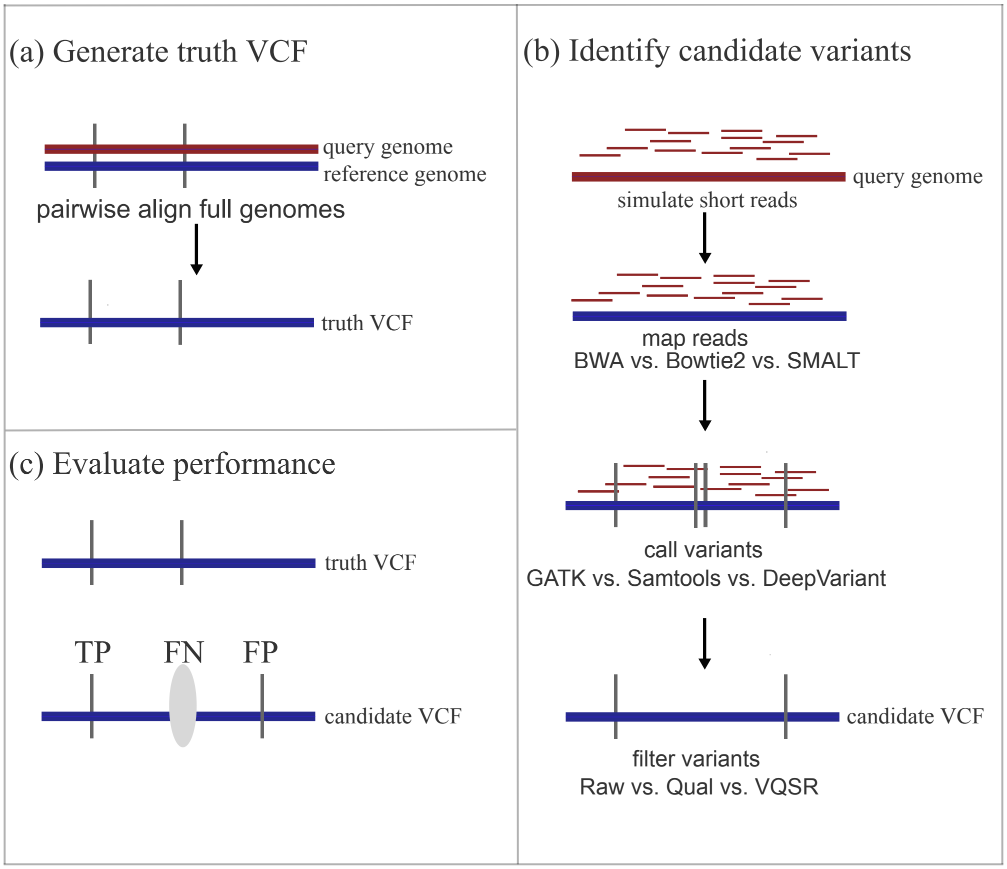


**Figure S1. Simulation approach to measure performance of variant calling tool combinations in recovering genome-wide variants.** (a) To generate truth sets of variants for query genomes (red) with respect to 12 *M. tuberculosis* different reference genomes (blue), we pairwise aligned the query genome (CDC1551) to each reference genome with MUMmer[43] (*nucmer* with the *maxmatch* option) and identified true variants in the query genome with respect to the reference genome *in silico*. (b) We then simulated Illumina sequence reads from the query genome. For each set of sequence data, we mapped reads with three commonly used mapping algorithms to the reference genome, called variants with three variant callers, and applied variant filters. This resulted in 24 candidate variant sets for each set of query sequence data (we did not apply VQSR to DeepVariant calls). (c) We compared each candidate VCF to the truth VCF to evaluate the performance of each tool combination.

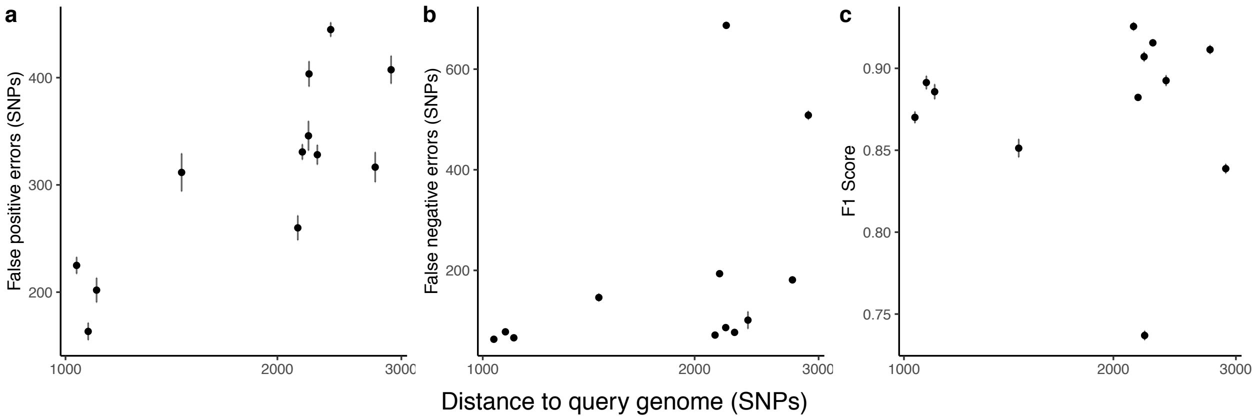


**Figure S2. Errors increase with increasing distance between query and reference genomes.** For a single tool combination (BWA/GATK), false positive (a) and false negative (b) errors increase with increasing distance between the reference and query genome, CDC1551. F1 score (c) also increases slightly with increasing distance between the reference and query genomes. Points and error bars indicate the mean and standard deviation false positive and false negative errors for 20 replicate sequence sets mapped against 12 reference genomes. The x-axis is log-transformed and y-axes are on different scales. Reference genomes are in Supplementary Table 2 and range from 1037 to 2901 SNPs distant to the query genome.

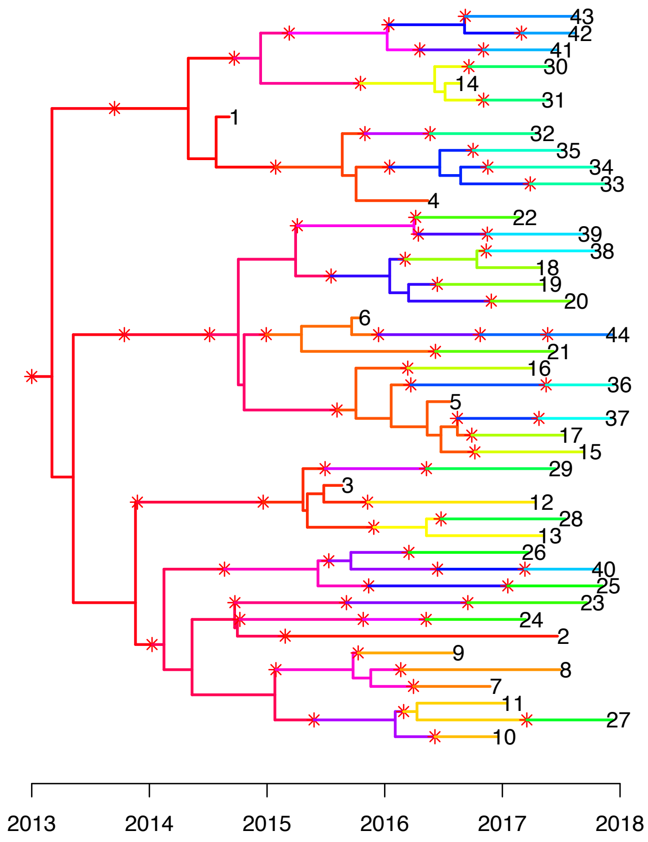


**Figure S3. Simulated *M. tuberculosis* outbreak transmission tree.** Transmission tree of a five-year clonal tuberculosis with a basic reproduction number, $R_0$, of 3 simulated with TransPhylo[4,5]. The tree topology represents the underlying *M. tuberculosis* outbreak phylogeny. Branches are colored according to infected hosts. Stars represent transmission events. Numbered tips represent sampled isolates.

**Supplementary Table 1.** *M. tuberculosis* variant calling pipelines compared in this study.

| Pipeline | Source | Quality control | Reference Genome | Mapper | Caller | Ploidy | Variants output | Variant filters | Regions excluded |
|---|---|---|---|---|---|---|---|---|---|
| A | | Read trimming with Sickle for reads with quality >20 and length > 30 | *M. tuberculosis* common ancestor | Bowtie 2 v. 2.2.9 | Samtools /bcftools v.1.3.1 | diploid | Single sample VCFs, variant sites only. | Mapping Quality > 30; depth ≥ 10X; fixed mutations (with a frequency of ≥ 75%); strand bias filter option on | Excluded all SNPs that were located in repetitive regions of the genome (for example, PPE/PEPGRS family genes, phage sequence, insertion or mobile genetic elements) that are difficult to characterize with short-read sequencing technologies; small insertions or deletions identified by VarScan (version 2.3.9) also excluded |
| B | https://github.com/TGenNorth/NASP | | 7199-99 NC_020089.1 | BWA | GATK | haploid | Multi-sample, variant sites only. | 10X threshold for depth of coverage and 0.9 proportion in base consensus | Masked any duplicated regions in the reference genome called by NUCmer |
| C | https://github.com/CPTR-ReSeqTB/UVP | FastQValidator, Kraken | H37Rv NC_000962.3 | BWA | GATK | diploid | Single sample VCFs, variant sites only. | Base quality >=20; mapping quality >=20, depth >=10X; <= 3 SNPs within 10 bp region. | Excluded repeat and problematic loci |
| D | | TrimGalore for reads with quality > 15 and stringency of 7 | H37Rv NC_000962.2 | BWA | Samtools /bcftools v.1.5 | haploid | Single sample VCFs, variant sites only. | Filtered SNPs within 15-bp of indels; QUAL > 100; (DP4[2]+DP4[3])/(DP4[0]+DP4[1]+DP4[2]+DP4[3]) > 0.75 | Excluded PE/PPE and other highly repetitive regions, plus 50 bp upstream and downstream of the same |
| E | 6 | | H37Rv NC_000962.2 | SARUMAN | Custom Perl script | haploid | Multi-sample FASTA, variant sites only. | Depth ≥ 10X, MAF ≥ 80% | Excluded 15 SNPs in repetitive regions such as PPE, PE_PGRS, ESX gene families that were false positives in Sanger sequencing |

**Supplementary Table 2. Summary of SNP variants identified by each pipeline.** Pipeline, number of samples passing quality filters, total internal SNPs included in FASTA, sensitivity to Sanger-sequence confirmed SNPs, mean and median pairwise distances, percentage of identical pairwise comparisons, percentage of sequence pairs falling within the 5 and 12-SNP thresholds for potential transmission. Internal SNPs refer to the number of SNPs in the VCF file that are variable within the outbreak.

| Pipeline | Samples | SNPs | Sensitivity | Mean pairwise | Median pairwise | Identical (%) | <= 5 SNPs (%) | <= 12 SNPs (%) |
|---|---|---|---|---|---|---|---|---|
| A | 86 | 94 | 92.9 | 3.1 | 2 | 11.1 | 80.7 | 99.9 |
| B | 86 | 68 | 72.9 | 3.8 | 1 | 29.7 | 69.8 | 96.5 |
| C | 68 | 352 | 88.2 | 45.5 | 40 | 0.0 | 0.0 | 0.2 |
| D | 86 | 416 | 90.6 | 54.0 | 42 | 0.0 | 0.4 | 8.0 |
| E | 86 | 85 | 100.0 | 5.5 | 4 | 9.6 | 63.1 | 89.2 |

**Supplementary Table 3. Comparison of pipelines for identifying SNP variants in the CDC1551 genome using the H37Rv reference genome.** Mapper, caller, filter, total true SNPs; SNPs reported by pipeline; TP, true positive; FN, false negative; FP, false positive; F1 score (harmonic mean of precision and recall); recall; and precision for each tool combination investigated. Filters include Raw, no filtering; Qual, filtering variants with a quality score of less than 40; and VQSR, filtering with GATK's Variant Quality Score Recalibration. Mean for 20 replicate sequence sets.

| Mapper | Caller | Filter | Truth | Reported | TP | FN | FP | F1 | Recall | Precision |
|---|---|---|---|---|---|---|---|---|---|---|
| Bowtie 2 | DeepVariant | Raw | 1107 | 1537.3 | 1046.2 | 60.8 | 491.1 | 0.791 | 0.945 | 0.681 |
| Bowtie 2 | DeepVariant | Qual | 1107 | 1071.5 | 991.4 | 115.6 | 80.2 | 0.910 | 0.896 | 0.925 |
| Bowtie 2 | GATK | Raw | 1107 | 1253.0 | 1043.0 | 64.0 | 213.9 | 0.882 | 0.942 | 0.829 |
| Bowtie 2 | GATK | Qual | 1107 | 1250.0 | 1039.0 | 68.0 | 211.0 | 0.882 | 0.939 | 0.831 |
| Bowtie 2 | GATK | VQSR | 1107 | 1033.8 | 992.9 | 114.1 | 41.0 | 0.928 | 0.897 | 0.961 |
| Bowtie 2 | Samtools | Raw | 1107 | 1425.0 | 1046.1 | 60.9 | 378.9 | 0.826 | 0.945 | 0.734 |
| Bowtie 2 | Samtools | Qual | 1107 | 1346.9 | 1040.3 | 66.7 | 306.6 | 0.848 | 0.940 | 0.772 |
| Bowtie 2 | Samtools | VQSR | 1107 | 1271.5 | 1034.8 | 72.2 | 236.7 | 0.870 | 0.935 | 0.815 |
| BWA | DeepVariant | Raw | 1107 | 1254.7 | 1045.7 | 61.4 | 209.1 | 0.886 | 0.945 | 0.833 |
| BWA | DeepVariant | Qual | 1107 | 1084.0 | 1009.5 | 97.5 | 74.5 | 0.922 | 0.912 | 0.931 |
| BWA | GATK | Raw | 1107 | 1238.9 | 1041.0 | 66.0 | 201.9 | 0.886 | 0.940 | 0.837 |
| BWA | GATK | Qual | 1107 | 1236.8 | 1036.7 | 70.3 | 200.1 | 0.885 | 0.936 | 0.838 |
| BWA | GATK | VQSR | 1107 | 1062.1 | 1018.9 | 88.2 | 43.2 | 0.939 | 0.920 | 0.959 |
| BWA | Samtools | Raw | 1107 | 1273.2 | 1040.3 | 66.7 | 232.9 | 0.874 | 0.940 | 0.817 |
| BWA | Samtools | Qual | 1107 | 1248.5 | 1037.8 | 69.2 | 210.8 | 0.881 | 0.937 | 0.831 |
| BWA | Samtools | VQSR | 1107 | 1180.2 | 1034.8 | 72.2 | 145.3 | 0.905 | 0.935 | 0.877 |
| SMALT | DeepVariant | Raw | 1107 | 1349.2 | 1048.5 | 58.5 | 300.6 | 0.854 | 0.947 | 0.777 |
| SMALT | DeepVariant | Qual | 1107 | 1095.9 | 1007.3 | 99.7 | 88.6 | 0.915 | 0.910 | 0.919 |
| SMALT | GATK | Raw | 1107 | 1277.7 | 1042.5 | 64.5 | 239.2 | 0.873 | 0.942 | 0.813 |
| SMALT | GATK | Qual | 1107 | 1276.2 | 1038.3 | 68.7 | 237.9 | 0.871 | 0.938 | 0.814 |
| SMALT | GATK | VQSR | 1107 | 1054.9 | 999.4 | 107.7 | 55.5 | 0.924 | 0.903 | 0.947 |
| SMALT | Samtools | Raw | 1107 | 1352.8 | 1039.2 | 67.8 | 313.6 | 0.845 | 0.939 | 0.768 |
| SMALT | Samtools | Qual | 1107 | 1318.0 | 1037.5 | 69.5 | 280.5 | 0.856 | 0.937 | 0.787 |
| SMALT | Samtools | VQSR | 1107 | 1251.8 | 1033.3 | 73.7 | 218.4 | 0.876 | 0.933 | 0.826 |

**Supplementary Table 4. Query genome and reference genomes used for mapping.**

| Strain | GenBank assembly accession | Lineage | Distance (SNPs) to Query Genome, CDC1551 |
|---|---|---|---|
| CDC1551 | GCA_000008585.1 | 4 | 0 |
| BCG Pasteur 1173P2 | GCA_000009445.1 | *M. bovis* | 2381 |
| F11 | GCA_000016925.1 | 4 | 1037 |
| KZN_1435 | GCA_000023625.1 | 4 | 1077 |
| Beijing_NITR203 | GCA_000364825.1 | 2 | 2901 |
| H37Rv | GCA_000195955.2 | 4 | 1107 |
| HN-024 | GCA_002356255.1 | 1 | 2170 |
| 18b | GCA_000835125.1 | 2 | 1462 |
| GM041182 | GCA_000253355.1 | *M. africanum* | 2279 |
| 96121 | GCA_000756545.1 | 1 | 2214 |
| EAI5/NITR206 | GCA_000389945.1 | 1 | 2754 |
| EAI5 | GCA_000422125.1 | 1 | 2138 |
| W-148 | GCA_000193185.2 | 2 | 2218 |

**Supplementary Table 5. Variant calling tools and versions used in the comparison of variant calling tools.**

| Software | Source | Version | Use |
|---|---|---|---|
| Bowtie2 | http://bowtie-bio.sourceforge.net/bowtie2/index.shtml | 2.3.4.2 | Maps reads |
| BWA | http://bio-bwa.sourceforge.net/ | 0.7.15-r1140 | Maps reads |
| SMALT | https://www.sanger.ac.uk/science/tools/smalt-0 | 0.7.6 | Maps reads |
| Sambamba | https://lomereiter.github.io/sambamba/ | 0.6.6 | Post-process BAM files, mark duplicates |
| Bcftools/Samtools | https://samtools.github.io/bcftools/ | 1.9-105-gaf6f0c9 | Variant calling/VCF processing |
| GATK | https://software.broadinstitute.org/gatk/ | 4.1.0.0 | Variant calling |
| DeepVariant | https://github.com/google/deepvariant | 0.7.0 | Variant calling |
| hap.py | https://github.com/Illumina/hap.py | 3.10 | Measures performance |
| MUMmer | http://mummer.sourceforge.net/ | 3.1 | Pairwise alignment |
| RAxML-ng | https://github.com/amkozlov/raxml-ng | 0.8.1 | Fit maximum likelihood trees |
| ART | https://www.niehs.nih.gov/research/resources/software/biostatistics/art/index.cfm | 2.5.8 | Sequence read simulation |
